# Crystal structures of the outer membrane transporter FoxA provide novel insights into TonB-mediated siderophore uptake and signalling

**DOI:** 10.1101/621219

**Authors:** Inokentijs Josts, Katharina Veith, Henning Tidow

**Author notes:** Corresponding authors: Inokentijs Josts, University of Hamburg, Department of Chemistry, Institute for Biochemistry and Molecular Biology, Martin-Luther-King-Platz 6, D-20146 Hamburg, Germany, Henning Tidow, University of Hamburg, Department of Chemistry, Institute for Biochemistry and Molecular Biology, Martin-Luther-King-Platz 6, D-20146 Hamburg, Germany.

## Abstract

Many microbes and fungi acquire the essential ion Fe^3+^ through the synthesis and secretion of high-affinity chelators termed siderophores. In Gram-negative bacteria, these ferric-siderophore complexes are actively taken up using highly specific TonB-dependent transporters (TBDTs) located in the outer bacterial membrane (OM). However, the detailed mechanism of how the inner-membrane protein TonB connects to the transporters in the OM as well as the interplay between siderophore- and TonB-binding to the transporter is still poorly understood. Here, we present three crystal structures of the TBDT FoxA from *Pseudomonas aeruginosa* (containing a signalling domain) in complex with the siderophore ferrioxamine B and TonB and combine them with a detailed analysis of binding constants. The structures show that both siderophore and TonB-binding is required to form a translocation-competent state of the FoxA transporter in a two-step TonB-binding mechanism. The complex structure also indicates how TonB-binding influences the orientation of the signalling domain.

## Introduction

Iron is one of the most abundant elements on earth and is essential for life. However, the bioavailability of iron in the environment is extremely low, and under aerobic conditions iron is found mostly as insoluble hydroxides. The demand for the ionic form of iron by all microorganisms growing in iron-limited conditions has led to the evolution of several efficient iron scavenging strategies. One of the predominant mechanisms by which microbes and fungi acquire iron is through the synthesis and secretion of small, specific high-affinity chelators termed siderophores, which keep iron in a chelated, soluble state (*1*). In Gram-negative bacteria, these ferric-siderophore complexes are actively taken up into cells using highly specific TonB-dependent transporters (TBDTs) situated in the outer bacterial membrane (OM) as well as specific transporters present in the bacterial inner membrane (IM) (*2*). The energy for this uptake process is derived from the proton motive force and relies on the energizing complex consisting of TonB/ExbB/ExbD situated in the bacterial IM (*3, 4*). TonB acts as a physical link between the transporters in the OM and the energizing complex in the IM (*5*). Siderophore binding facilitates TBDT contact with the C-terminus of TonB through an allosteric mechanism, which exposes the TonB-binding site within TBDT, known as the TonB box, to the periplasm (*6*). Association of TBDTs with TonB establishes the main point of contact with ExbB/ExbD and the proton motive force provides the energy needed for the translocation of siderophores through the lumen of the OM barrel. The precise mechanism of this translocation process is not yet fully understood but is thought to involve a mechanical extraction or unfolding of the plug region from within the barrel lumen (*7*). Furthermore, a subclass of TBDTs possesses an N-terminal signalling domain, which regulates gene transcription of target operons, often participating in siderophore uptake and processing (*8*). The activation of these signalling cascades is both ligand- and TonB-dependent, however the molecular details of this signalling process and its activation remain highly elusive (*9, 10*). It is speculated that the N-terminal pocket of the signalling domain is involved in the interactions with the σ-factor regulator proteins. To date, the crystal structures of the intact TBDT FpvA with the fully-resolved signalling domain suggest that there is a high degree of flexibility at the N-terminal region of the transporter (*11*). However the putative site involved in contacting the regulator protein is tucked away beneath the barrel lumen.

*Pseudomonas aeruginosa* is a Gram-negative bacterium and an opportunistic human pathogen, which is a major cause of hospital-acquired infections in immunocompromised patients. In patients with cystic fibrosis *P. aeruginosa* lung infections are usually associated with increased mortality rates. *P. aeruginosa* is able to utilise a range of xenosiderophores, i.e siderophores produced by other bacteria or fungi in order to scavenge free iron. Such instances of so-called ‘siderophore piracy’ highlight bacterial adaptability and potential for colonising in an extremely broad range of environmental niches. One example of siderophore piracy is the uptake of ferrioxamine B, a hydroxamate siderophore produced by many *Streptomyces* species, by the specific OM transporter FoxA (*12*). FoxA belongs to the family of TBDTs (transducers) and is involved in ferrioxamine B transport as well as modulation of transcriptional cascades in the bacterial cell. Ferrioxamine B uptake comes at a relatively low energetic cost, when compared with the production of native siderophores such as pyoverdin and pyochelin (*13*). Indeed, when grown in the proximity of *Streptomyces ambofacients*, several *Pseudomonas* species do not produce their own siderophores and instead parasitize on the siderophores of their neighbour by expressing the ferrioxamine B transporter, FoxA (*14*). Here, we determined several crystal structures of FoxA from *P. aeruginosa*, in the apo state as well as in complex with the siderophore ferrioxamine B and TonB. Using a hybrid approach combining X-ray crystallography with biophysical interaction studies, we provide novel insights into TonB-mediated siderophore uptake across the bacterial OM as well as TonB-dependent signalling. Our results indicate that the transporter can exist in several different conformations, and that both substrate- and TonB-binding is required to form a translocation-competent state of the FoxA transporter in a two-step TonB-binding mechanism necessary for transport function.

## Results and Discussion

### Ternary structure of FoxA bound to ferrioxamine B and TonB_Ct_

Active uptake of siderophores such as ferrioxamine B across the outer membrane (OM) relies on the establishment of a physical contact between the specific transporter and inner membrane (IM)-tethered TonB. In order understand the mechanism of complex formation between FoxA and TonB we determined the crystal structure of the ternary complex consisting of FoxA bound to ferrioxamine B and the C-terminal TonB domain (residues 251-340, referred to hereafter as TonB_Ct_). Two complexes were present in the asymmetric unit, with crystal contacts generated through the exposed soluble regions of both proteins (Figure S1A,B). We could resolve almost an entire FoxA molecule including the signalling domain (residues 53-820, with 11 amino acids missing from the N-terminus after the signal peptide) as well as the full TonB_Ct_. We can identify two main modes of contact between FoxA and TonB_Ct_. Similar to the FhuA and BtuB-TonB complexes (*15, 16*) the primary interaction site between FoxA and TonB_Ct_ occurs through β-augmentation with parallel strands forming between residues 332-337 of TonB and residues 142-146 of FoxA (Figure 1A,B). In addition to the observed β-augmentation, the unstructured polypeptide segment upstream of the TonB box (residues 135-141) is forms complementary contacts with the surface of TonB molecule mediated by backbone H-bond interactions as well as side-chains Leu_137_, Met_139_ and Val_142_ located in a small cavity on the surface of TonB (Figure 1C). Most of the TBDTs do no harbour this extra binding motif upstream of the TonB box since it would only be present in TBDTs involved in regulating signalling events at the OM via the additional N-terminal domain in the periplasm. The secondary point of contact involves the side chains of residues in the periplasmic loops and barrel interior of FoxA with several residues in the loops of TonB facing the barrel. These contacts are mediated by electrostatic forces between TonB_R271_ and FoxA_D352_, and between TonB_R266_ and FoxA_D355_ as well as a side-chain (FoxA_E316_) to the backbone carbonyl (TonB_S300_) via a hydrogen bond (Figure S2). These contacts provide a secondary site of attachment for the TonB fragment and tether the C-terminal region of TonB to the barrel. This tethering locks the orientation of TonB against the barrel and the membrane plane. An overlay of the two complexes found in the asymmetric unit reveals that TonB_Ct_ and the signalling domain are rotated by 9° with respect to the barrel and the membrane plane (Figure S2). It is evident that TonB_Ct_ and the signalling domain experience some degree of flexibility in the distal part of the complex, whereas the proximal part of the complex is most likely stabilized by the contacts at the secondary site. In the previous structural models (*15, 16*), TonB sits in close proximity to the β-barrel, almost parallel to the putative lipid bilayer plane. In our crystal structure, the TonB fragment is located almost perpendicular to the β-barrel and the membrane plane highlighting the potentially dynamic nature of TBDR-TonB complexes, which has been suggested by recent EPR experiments (*17*) (Figure S3). Overall, the structure of the ternary FoxA-ferrioxamineB-TonB_Ct_ complex presented in this work reveals both the structure of the N-terminal signalling domain as well as a markedly different orientation of the TonB_Ct_ relative to the TBDT compared to previously determined ternary structures of FhuA and BtuB (*15, 16*).

**Figure 1.**
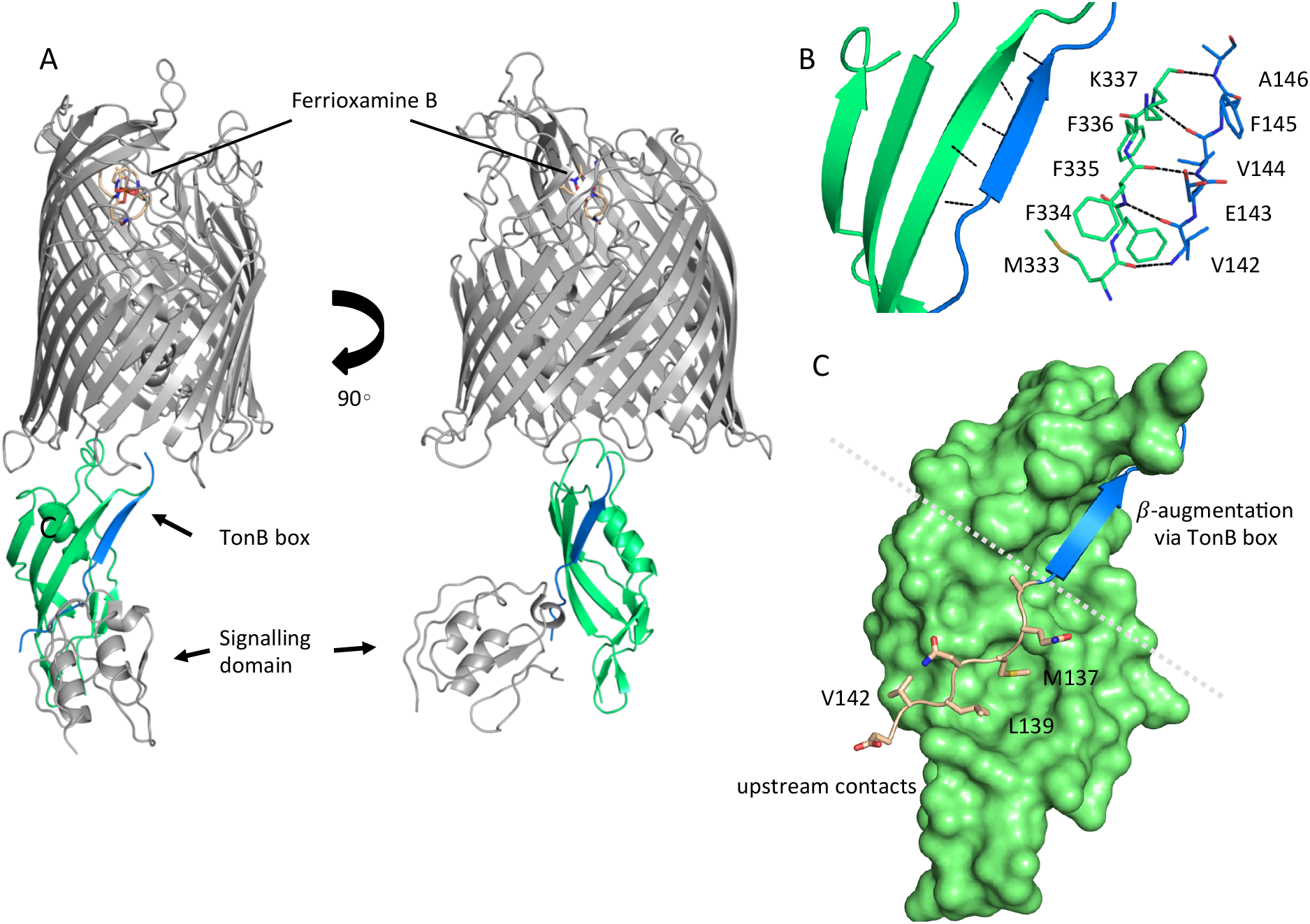
Complex formation between ferrioxamine B-bound FoxA and TonB is driven by multiple binding sites. A) Overview of the ferrioxamine B-bound FoxA-TonB_Ct_ complex. TonB_Ct_ (green) interacts with the TonB box (blue) of FoxA (grey). B) Parallel β–strands formed between the TonB box of FoxA (blue) and TonB_Ct_ (green) through β-augmentation. All contacts are mediated predominantly by backbone hydrogen bonds between the two proteins. C) Polypeptide stretch (pink) upstream of the TonB box (blue) forms additional contacts with the surface of TonB (green).

To understand the conformational changes occurring in FoxA in response to ferrioxamine B and TonB_Ct_ we also determined the crystal structure of FoxA in its apo state (Figure 2A). In this structure a large solvent-exposed lumen faces the extracellular side of the membrane and is filled with solvent molecules. No electron density was present for the last two amino acids of the TonB box region and the N-terminal signalling domain of FoxA (residues 45-143), most likely due to the high flexibility of the linker connecting the plug domain and the signalling domain, as previously observed in the structures of FecA (*18, 19*) and FpvA (*11, 20, 21*). Inspection of both apo as well as ligand- and TonB_Ct_-bound crystal structures of FoxA revealed that in the apo state the TonB box is predominantly occluded in the interior of the barrel domain. A comparison of the plug domain conformations in both of our FoxA structures indicates that the TonB box must be displaced by approximately 22 Å from the folded plug domain in order to allow for β-augmentation to occur with TonB (Figure 2B).

**Figure 2.**
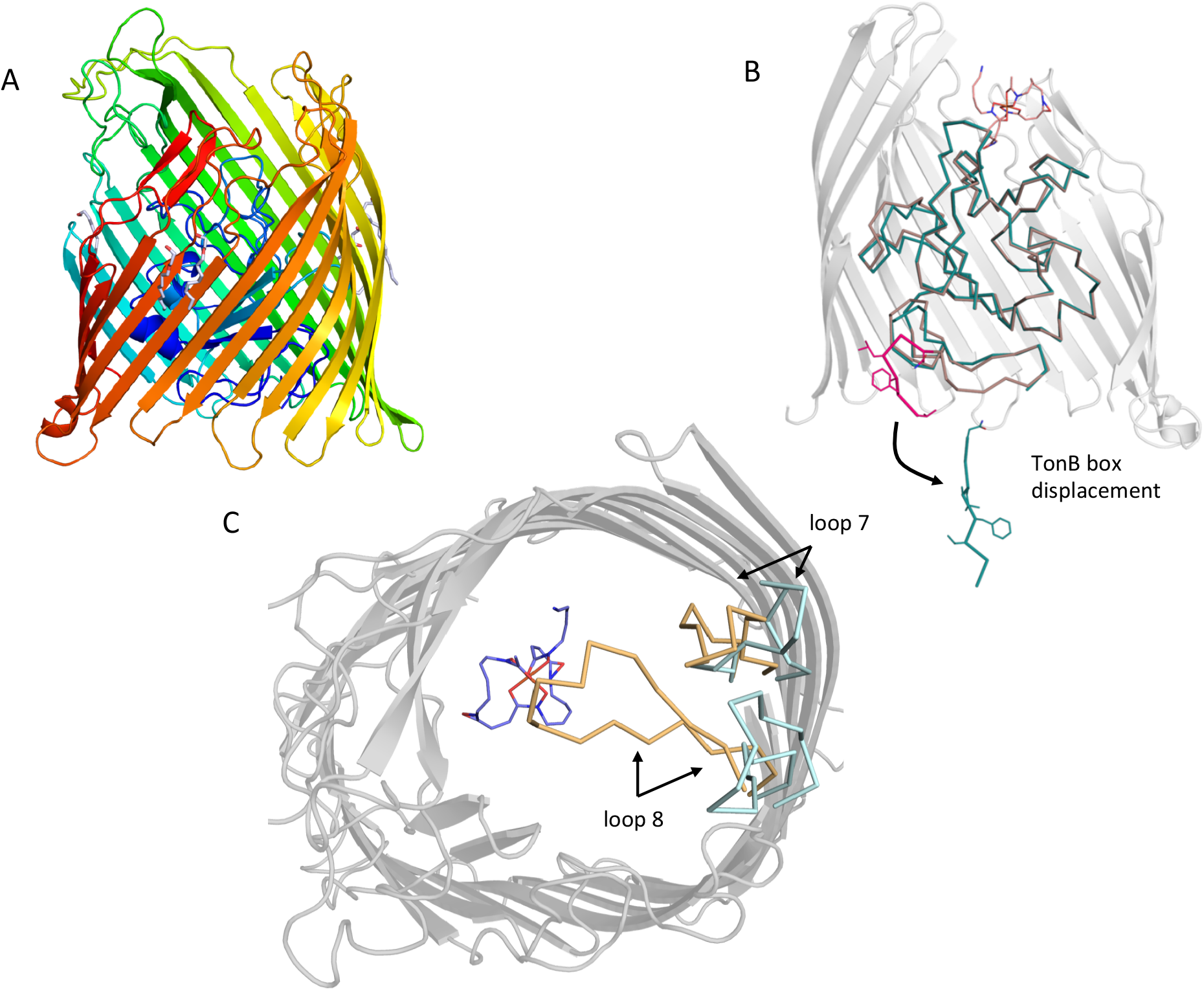
Conformational changes in the plug domain and extracellular loops of FoxA in response to ferrioxamine B and TonB binding. A) The overall fold of apo FoxA consists of a 22-stranded β-barrel lined by the small globular plug domain within the lumen. The structure is color-coded from blue (N-terminus) to red (C-terminus). B) Structural rearrangements within the plug domain necessary to accommodate interactions with TonB_Ct_. The region of the polypeptide being part of the TonB box common to both FoxA structures is highlighted in pink. This region is displaced by approximately 22 Å into the periplasm. Slight conformational changes are also observed throughout the rest of the plug domain (blue: ternary complex / brown: apo state). C) Loops 7 and 8 enclose the bound siderophore within the hydrophobic cavity to prevent its dissociation and reduce permeation across the bacterial membrane during the process of siderophore uptake. Loop closure is only evident once the FoxA is bound with ferrioxamine B and the TonB_Ct_ fragment (loops coloured brown), indicative of allosteric communication between the extracellular and periplasmic regions of the transporter (blue loops correspond to the apo FoxA).

Compared to the apo FoxA structure the ternary complex of FoxA-FoaB-TonB_Ct_ also reveals substantial loop movements. Displacement of loops 7 and 8 by approx. 7 Å on the extracellular side of the membrane leads to the closure of the barrel lumen on both sides of the membrane, limiting the access to the barrel lumen (Figure 2C). Mechanistically, this would prevent the dissociation of the siderophore during translocation and opening of the entry channel within the barrel.

### Biophysical characterization of the interactions between FoxA and TonB_Ct_

Previous structural and biochemical investigations into the mechanisms of TBDT activation and substrate uptake have shown that siderophore binding usually leads to an unwinding of either an N-terminal helix or a stretch of polypeptide within the plug domain bearing the TonB box motif. This mechanism of polypeptide unwinding, initiated by concerted small motions throughout the plug, allows the C-terminus of TonB to make contact with the loaded transporter molecule. Moreover, insights into TBDT association with TonB paint a very complex, and often conflicting picture of substrate-dependent transporter activation and TonB binding. One model suggests a constitutively bound TonB-TBDT complex (*22, 23*), whilst another proposes a cooperative mode of transporter-TonB interactions that is driven by initial substrate capture by the TBDT (*24, 25*).

Therefore, we sought to understand the nature of FoxA-TonB interactions using isothermal titration calorimetry (ITC) to characterize the thermodynamics of FoxA-TonB interactions. For this purpose, we have reconstituted FoxA into MSP1D1 nanodiscs in order to minimize the detergent mismatch (*26, 27*). Additionally, nanodiscs (ND) provide a lipidic scaffold and alleviate the deleterious effects detergents might have on the conformation of the transporter. Titration of TonB_Ct_ into apo FoxA-ND complexes resulted in strong, saturable exothermic heats indicative of protein association. ITC data were fitted to a single binding site model and yielded a K_d_ of 107 ± 30 nM. The binding process is enthalpically driven with a large, negative ΔH of −11.01 ± 0.3 kcal mol^−1^ and entropically unfavourable with a negative TΔS value of −1.47 kcal mol^−1^ (Figure 3A). No binding was observed when TonB_Ct_ was titrated into empty nanodiscs. The observation of tight binding between TonB_Ct_ and apo FoxA in nanodiscs is in contrast to similar experiments performed with FhuA and TonB_Ct,_ for which no binding could be observed (*27*). We speculate that the presence of the N-terminal domain and the interacting region upstream of the TonB box in FoxA is responsible for the differences in the binding modes between these two transporters.

**Figure 3.**
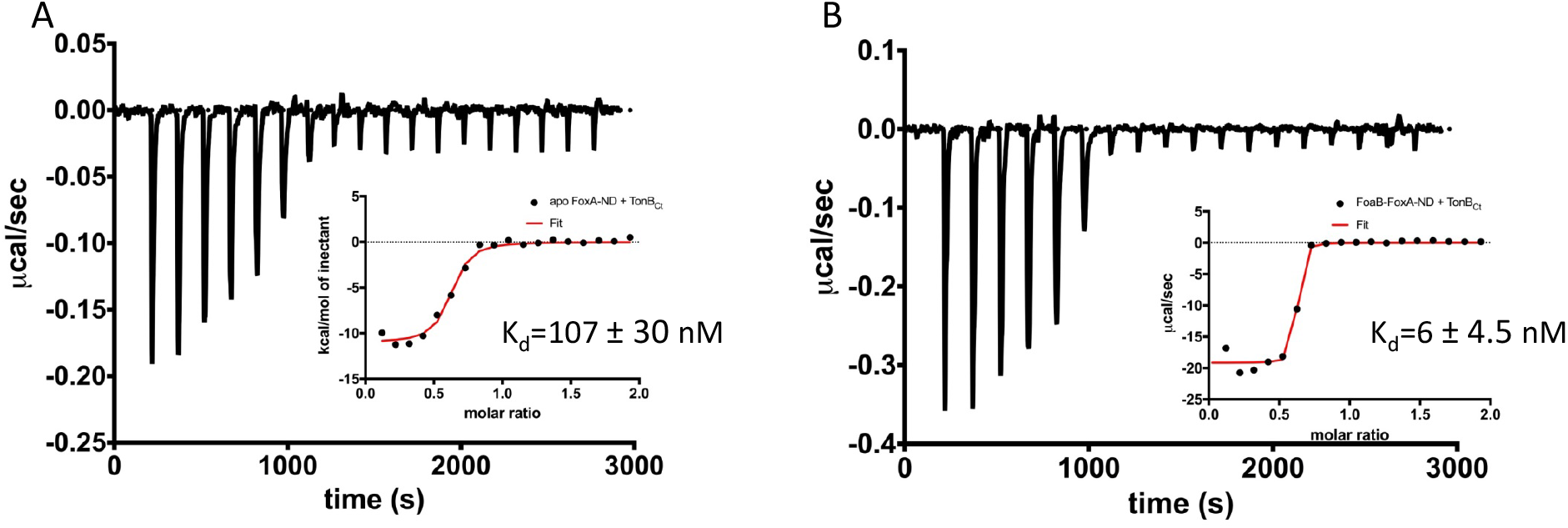
Two distinct modes of association between FoxA and TonB as revealed by the thermodynamics of complex formation. ITC profiles showing titration of TonB_Ct_ (150 μM) into 15 μM apo FoxA nanodisc complexes (A) and 15 μM ferrioxamine B-FoxA nanodisc complexes (B). Insets show the integrated heats of binding with a single-site fit to the data.

Next, we analyzed the interactions between TonB_Ct_ and ferrioxamine B-bound FoxA in lipid nanodiscs. The data were also fitted to a single binding site model. Our ITC experiments showed that in the presence of ferrioxamine B the binding affinity is increased 17-fold yielding a K_d_ value of 6 ± 4.5 nM. The thermodynamic parameters of the association reaction also differ such that the ΔH value decreases drastically to −19.16 ± 0.4 kcal mol^−1^ and the entropic contribution becomes even more unfavourable with TΔS of −7.86 kcal mol^−1^ (Figure 3B).

We speculate that the decrease in the entropy of TonB_Ct_ binding in the presence of ferrioxamine B most likely arises from large conformational restrictions of the flexible and highly mobile TonB and signalling domains as well as folding/desolvation events involved in association with TonB_Ct_. The difference in thermodynamics between the two binding reactions also suggests several distinct binding modes between TonB_Ct_ and FoxA, which rely on the siderophore capture. The negative entropy is compensated for by a large decrease in the enthalpy of binding, which is driving the association reaction. Our ITC data suggest that in the presence of ferrioxamine B, FoxA is able to form a much tighter complex with TonB_Ct_. Since β-augmentation between the TonB box of FoxA and TonB_Ct_ is driven predominantly by hydrogen-bonded interactions, the stark decrease in the enthalpy of binding in our ITC could be explained by the formation of these additional contacts. The increased affinity and drastically reduced enthalpy of association is indicative of a much larger surface area participating in the complex formation process compared with the thermodynamics of apo FoxA-TonB_Ct_ complexes. Altogether, the interaction studies presented here strongly support a cooperative mechanism of siderophore-dependent TonB capture by FoxA and that two distinct TonB-binding events can occur at the FoxA transporter.

To delineate the interactions between FoxA and TonB we generated FoxA variants with truncations in the signalling domain and analysed complex formation using analytical size-exclusion chromatography. Full-length FoxA exhibits three distinct elution profiles corresponding to apo protein, FoxA-TonB_Ct_ complex and the ternary complex FoxA-ferrioxamine B-TonB_Ct_ confirming our ITC findings of two distinct TonB_Ct_-bound states, with and without the siderophore (Figure S4A). Deletion of residues 64-130, which correspond to the majority of the signalling domain, yet retaining the sequence upstream of the TonB box observed in our crystal structure, had no effect on the constitutive and cooperative binding of TonB_Ct_ to FoxA (Figure S4B). However, the deletion of residues 64-143, which also include the upstream binding motif of the TonB box, abrogated the constitutive binding of TonB_Ct_ to FoxA, whilst retaining the ability to cooperatively form the ternary complex between FoxA, ferrioxamine B and TonB_Ct_ (Figure S4C). In combination with our structure of the ternary complex, we propose a two-step binding mechanism: The constitutive mode of TonB_Ct_ binding is mediated by the stretch of amino acids located upstream of the TonB box, and the interaction with ferrioxamine B would result in the allosteric release of the TonB box from within the barrel interior allowing the formation of a very tight complex between ferrioxamine B-bound FoxA and TonB, which is necessary for downstream translocation events leading to siderophore uptake.

### Loop movements establish additional contacts with the bound siderophore

Siderophore capture by a specific TBDT is an integral part of establishing the necessary contacts with the TonB/ExbB/ExbD complex, which provides the energy for substrate translocation. As we have demonstrated, the full engagement of TonB by FoxA is dependent on the presence of ferrioxamine B. To understand how FoxA interacts with its substrate ferrioxamine B, we co-crystallised and determined the structure of FoxA with ferrioxamine B in absence of TonB_Ct_ and compared this structure with apo FoxA as well as the ternary complex. Our ferrioxamine B–bound structures of FoxA revealed a common siderophore binding mode similar to other TBDTs. The electron density for the bound ferrioxamine B in both structures allowed us to unambiguously model the conformation of the ligand as judged by the Polder omit maps. (Figure S5 A,B). Ferrioxamine B occupies a highly hydrophobic cavity and is stabilised predominantly through hydrophobic and van der Waal interactions via the aromatic residues found in the surrounding loops and the plug domain inside the cavity (Figure 4 A,B). Several hydrogen bonds are observed between the Tyr_805_ and FoaB_O23_, Tyr_218_ and FoaB_N12_, and His_374_ and FoaB_O22_. The His_374_ residue, located on the 3^rd^ extracellular loop, faces the octahedrally-coordinated Fe(III) and allows the imidazole side chain to form a hydrogen bond with the hydroxamate groups of the siderophore. Additionally, a number of hydrogen bonds are observed between ferrioxamine B and the surrounding water molecules. One of the water molecules involved in the coordination is stabilised through hydrogen bonds by Gln_441_ (Figure 4B).

**Figure 4.**
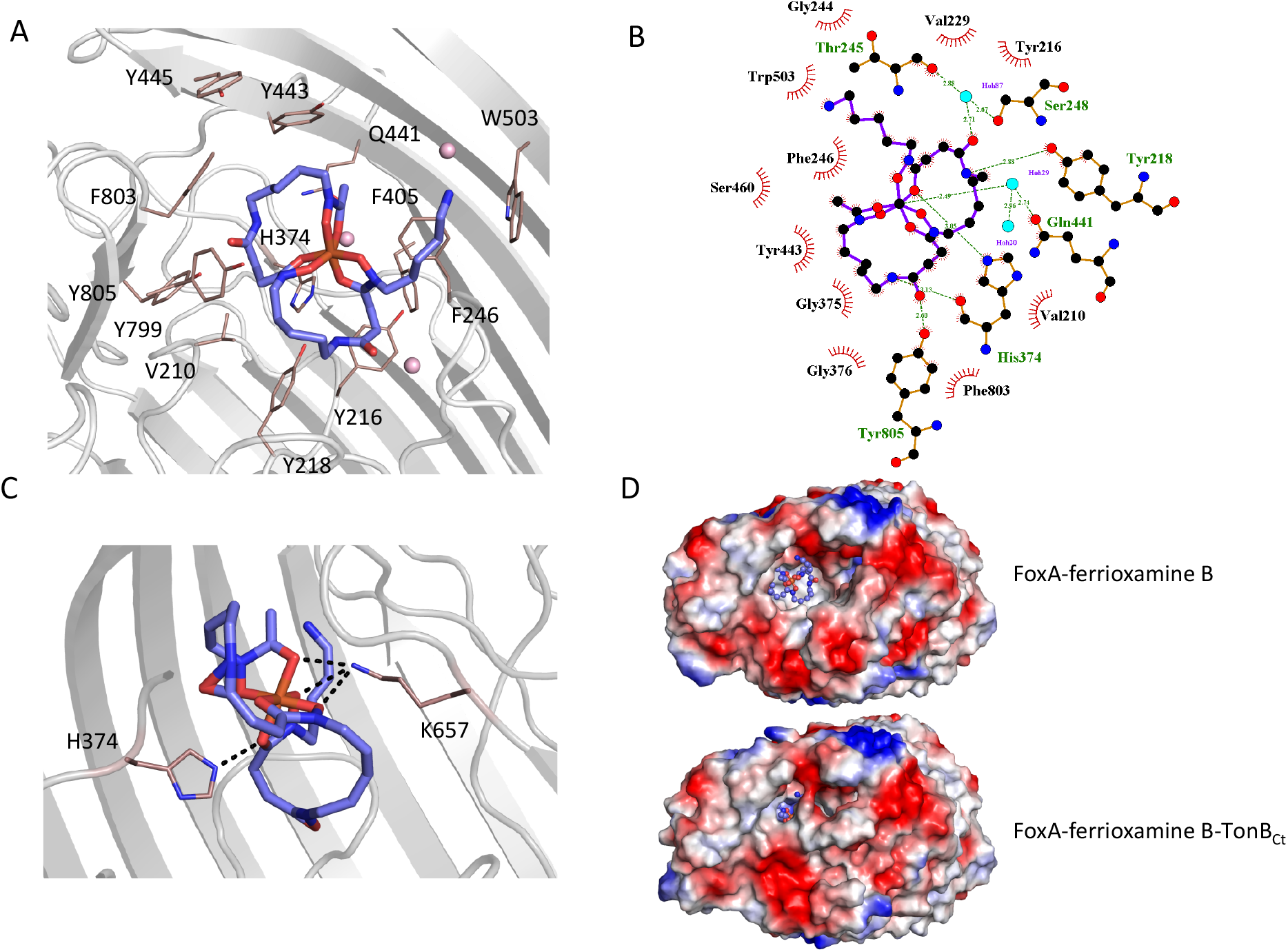
Ferrioxamine B interactions with FoxA reveal the basis for high-affinity siderophore capture. A) and B) Interaction of ferrioxamine B by hydrophobic and aromatic residues lining the binding pocket of FoxA. Several hydrogen bonds are also observed between ferrioxamine B and residues His_374_ and a Gln_441_-H_2_O network, respectively. C) The closure of loop 8 results in additional hydrogen bonds between ferrioxamine B and the ε-amino group of Lys_657_ facing the siderophore, which becomes locked from both sides by hydrogen bonds. D) Large hydrophobic cavity facing the extracellular milieu occupied by ferrioxamine B. In the apo/ferrioxamine B structures the cavity and ferrioxamine B are solvent exposed (top), whereas in the ternary complex (bottom) the loop closure sequesters ferrioxamine B inside the barrel.

**Figure 5.**
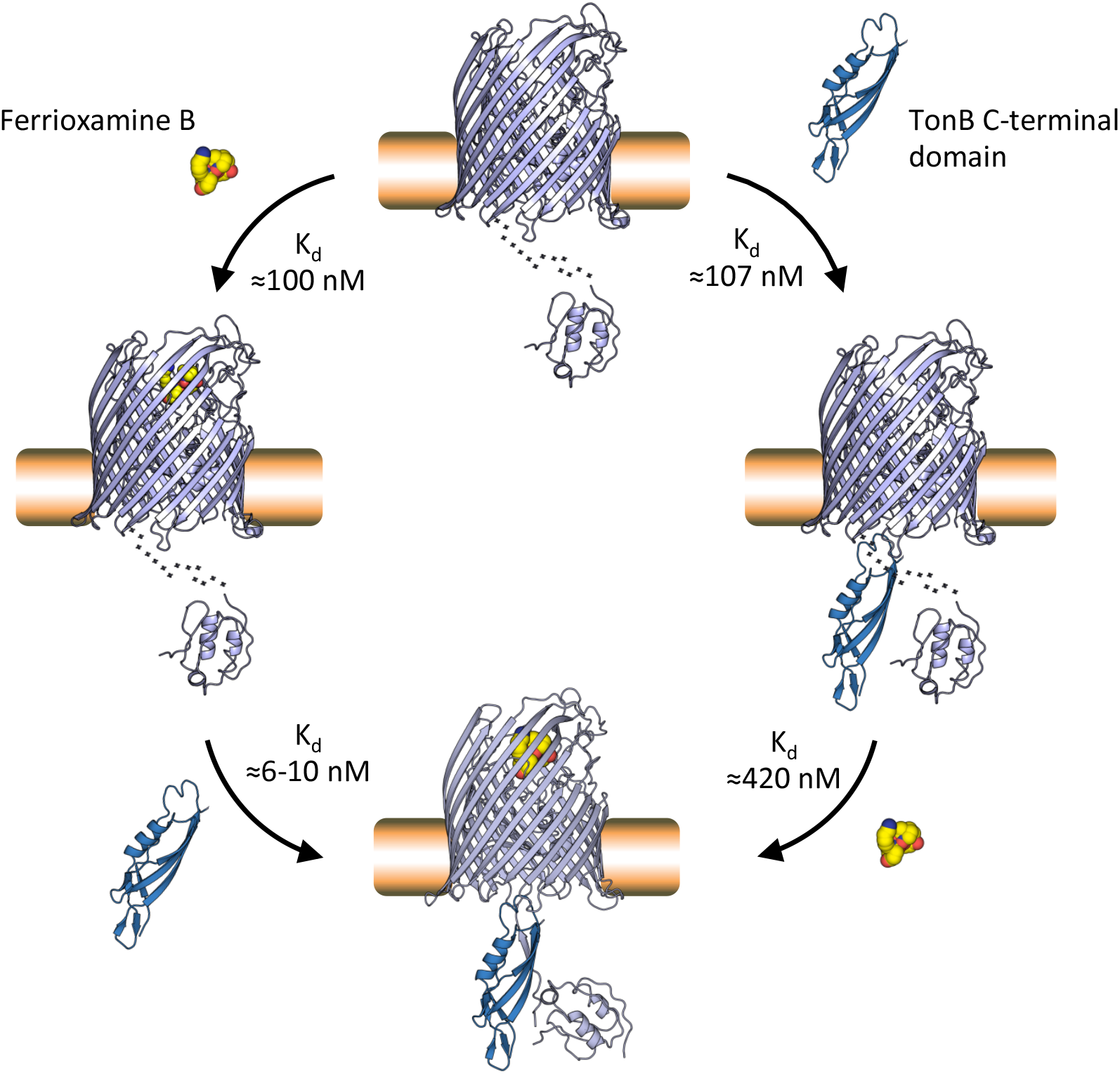
Proposed mechanism of TonB-mediated ferrioxamine B uptake via the FoxA transporter. Our studies suggest that FoxA can exist in several states depending on the abundance of ferrioxamine B in the environment and the occupancy of TonB by other TBDRs. FoxA is able to interact with ferrioxamine B or engage with TonB with near-equal affinity (K_d_≈100 nM) in a constitutive fashion. The presence of ferrioxamine B, however, is necessary for the expulsion of the TonB box into the periplasm and the formation of the full, translocation-competent ternary complex through β-augmentation (K_d_≈6-10 nM). This very high-affinity interaction provides the necessary contacts in the complex for subsequent steps of the plug domain re-modelling or expulsion, necessary for siderophore translocation through the lumen of the barrel.

The inward movement of loop 8, which is seen in the ternary complex with TonB_Ct_, places Lys_657_ into close proximity of the bound ferrioxamine B with its ε-amino group protruding towards the hydroxamate groups coordinating Fe^3+^ on the opposite side of His_374_ (Figure 4C). These interactions presumably enforce the directionality of siderophore passage in the instance where the high-affinity site on the plug domain is modified during the partial unfolding and provides additional stabilising contacts to the siderophore prior to steps leading to its transport through the lumen of the barrel.

We measured the interaction between ferrioxamine B and purified FoxA using tryptophan fluorescence quenching experiments and ITC. Titration of ferrioxamine B to FoxA purified in nonyl glucopyranoside lead to concentration-dependent quenching of Trp fluorescence and allowed us to calculate the dissociation constant (K_d_) of 100 nM, with a 1:1 stoichiometry (Figure S5C). Our ITC experiments titrating ferrioxamine B into FoxA yielded a K_d_ value of 180 ± 140 nM, which agrees well with our fluorescence experiments (Figure S5D). Such strong association is also observed in other widely studied TBDT-siderophore complexes (*28*) and reflects the highly specific nature of these transporters at capturing extremely scarce siderophore-iron chelates from the extracellular milieu. Our crystal structures of apo FoxA and the ferrioxamine B-bound states revealed only minor conformational perturbations of the extracellular loops and the periplasmic side of the plug domain in FoxA upon binding with the siderophore. However, when ligand-loaded, FoxA association with TonB_Ct_ leads to the closure of the extracellular loops, shielding the bound siderophore inside the barrel. It is worth mentioning that the ability of FoxA to bind ferrioxamine B is unaffected by the bound TonB_Ct_ (Figure S6), meaning that the movement of the loops is not initiated by the binding of TonB to the apo state of the transporter.

### Conclusions

Our structures enable us to postulate a mechanistic model for the TonB-mediated uptake of ferrioxamine B by the TBDT FoxA. The FoxA transporter can form a constitutive complex with TonB, even in the absence of ferrioxamine B. Ferrioxamine B binding to this constitutive state of FoxA-TonB would then lead to the full, high-affinity contact with TonB via the formed β-augmentation and initiate a cascade of structural re-arrangements within the plug domain to allow for the passage of the substrate through the barrel lumen. Likewise, depending on the occupancy of TonB, FoxA can sequester free ferrioxamine B from solution and remain siderophore-bound until it is able to engage with TonB for transport of the siderophore inside the periplasm (Fig 5). Our structure also offers potential insight into signal transduction necessary for the regulation of transcription of the FoxA operon. The signalling domain, visible in the ternary complex, is exposed N-terminally to the periplasmic space, where is can at some stage of the transport engage with the sigma factor regulator protein FoxR situated in the IM. Such an orientation of the signalling domain differs starkly from previous structural studies involving FpvA (*11, 21*).

## Materials and Methods

### Materials

The detergents used for purification were from Anatrace (Maumee, OH, USA) or Glycon (Luckenwalde, Germany). Desferrioxamine B was purchased from Sigma-Aldrich. All other chemicals were of analytical grade and obtained from Roth (Karlsruhe, Germany) or Sigma Aldrich / Merck (Darmstadt, Germany).

### Protein expression and purification

Full-length FoxA gene from *Pseudomonas aeruginosa* stain PAO1 was cloned into a modified pET28a vector bearing a C-terminal TEV cleavage site prior a His_6_-tag. Protein overexpression was carried out in *Escherichia coli* Lemo21 cells (*29*) in 2xTY media supplemented with NPS (50 mM Na_2_HPO_4_, 50 mM KH_2_PO_4_, 25 mM NH_4_SO_4_) and 5052 mix (0.05% glucose, 0.2% lactose, 0.5% glycerol) and 0.5 mM L-rhamnose. Cells were grown to an OD_600_ of 1 at 37 °C, the temperature was reduced to 20 °C and 0.1 mM IPTG was added for further 16 hours. Cells were lysed in 30 mM Tris pH7.5, 200 mM NaCl, 10% glycerol using the high-pressure homogenizer (EmulsiFlex-C3, Avestin) and cell debris was removed by centrifugation at 22,000 g for 30 min. 1% Triton X-100 was added to the clarified cell lysate and incubated for 1 hr at 4 °C. The outer membrane fraction was isolated by a second centrifugation step at 100,000 g and the resuspended pellet was solubilised overnight in 1% octyl glucopyranoside (OG). Insoluble material was removed by another centrifugation step at 100,000 g for 20 min. Solubilised OM fraction was applied to the Ni-NTA resin followed by subsequent washes with buffer containing 25 mM imidazole and 0.4% nonyl glucopyranoside (NG) or 0.4% C8E4. Protein was eluted with 250 mM imidazole in buffer with 0.4% NG or C8E4 and tobacco etch virus protease (1:10 w/w) was added to the eluted fractions overnight. After reverse Ni-NTA purification, the protein was concentrated and passed over a Superdex S200 10/300 size exclusion column.

TonB_Ct_ (TonB1 from from *Pseudomonas aeruginosa* strain PAO1, residues 251-340) was overexpressed in *Escherichia coli* BL21 Gold cells, grown at 37 °C in LB medium. Cells were induced with 0.2 mM IPTG at OD_600_ of 0.6-0.8 and the temperature was reduced to 20 °C. After 12 hours cells were spun down and lysed in 30 mM Tris pH 7.5, 500 mM NaCl, 10% glycerol using the EmulsiFlex-C3 (Avestin) homogeniser and cell lysate was centrifuged at 40,000 g to remove the cell debris. Cleared cell lysate was supplemented with 20 mM imidazole and loaded onto the 5-ml HisTrap Ni-NTA column. After several washes the protein was eluted with resuspension buffer supplemented with 300 mM imidazole. TEV was added to the pooled fractions containing TonB_Ct_ and reverse-purification was performed the next day to remove the TEV and cleaved His_6_ tag. TonB_Ct_ was concentrated and stored at −80 ᵒC until further use.

MSP1D1 was expressed and purified as previously described (*30, 31*) (*32*). Briefly, MSP1D1 in pET28a vector was transformed in *E. coli* strain BL21 (DE3) and grown in *terrific broth* (TB) media at 37 °C. At an OD_600_ of 1.5 the protein expression was induced by adding 1 mM isopropyl ß-D-1-thiogalactopyranoside (IPTG) and cells were grown for 4 h at 37 °C. Cells were harvested by centrifugation at 3000 g, resuspended in lysis buffer (50 mM Tris pH 8.0, 500 mM NaCl) with 1% Triton X–100 and broken using sonication. The cleared lysate was loaded onto a HisTrap column and washed with ten column volumes each of lysis buffer containing 1% Triton X-100 and 50 mM cholate, respectively. MSP1D1 was eluted with buffer containing 500 mM imidazole, and fractions containing pure protein were pooled and incubated with TEV protease overnight. Subsequently, the protease and cleaved His-tag were separated by applying a second IMAC chromatography step and MSP1D1 without His-tag was concentrated up to 400 µM and stored at −80 °C until further use.

### Analytical size-exclusion chromatography (SEC)

Truncation mutants of FoxA (Δ64-130 and Δ64-143 residues) were generated using the standard QuikChange PCR mutagenesis protocols. Analytical SEC analyses of complex formation between FoxA, truncated FoxA and TonB_Ct_ were performed using a Superdex S200 10/300 column. The buffer in these experiments consisted of 25 mM HEPES pH7.4, 150 mM NaCl, 0.4% NG. In all cases, 15-25 μM FoxA or its truncated forms were mixed with 5-fold excess TonB_Ct_ and 10-fold excess ferrioxamine B (where appropriate) and injected onto the analytical SEC column.

### Crystallisation

Crystals of apo FoxA purified in C8E4 were grown by sitting drop vapour diffusion technique. 1 μl of purified protein (5-10 mg/ml) was mixed with 1 μl of 1.8-2 M ammonium sulfate solution and 0.1 M HEPES pH 7. Crystals appeared overnight and grew out of phase separation liquid clusters with crystal sizes, limited by the liquid phase, reaching approximately 30-70 μm. For crystallisation of FoxA with ferrioxamine B in NG detergent, the ligand was added in excess to the protein solution and incubated for at least 30 min on ice prior to setting up crystallisation trays. Crystals were cryo-protected by a step-wise addition of glycerol to a final concentration of 18-20% (v/v). For the crystallisation of the FoxA-FoaB-TonB_Ct_ complex, FoxA-FoaB was incubated with 2-fold excess TonB_Ct_, and passed through the size-exclusion column. Fractions corresponding to the complex were pooled and set up in crystallisation screens. The complex crystallised in the same condition as the apo FoxA and FoxA-FoaB, containing 1.8-2 M ammonium sulfate, 0.1 M HEPES pH 7. These crystals were cryo-protected by soaking the crystals in the crystallisation condition supplemented with 20% ethylene glycol for 2-3 min.

### Structure determination

All X-ray diffraction data was collected at 100 K. Data were collected at the PETRA III/EMBL P14, ESRF ID30B and BESSY 14.1 beamlines. All datasets were processed with XDS (*33*), and merged with AIMLESS (*34, 35*). All final data were merged from two individual datasets. Unit cell parameters and space groups are given in Table I. We used the FhuA model from *Escherichia coli* (PDB:1BY3) as a molecular replacement candidate (40-45% sequence identity to FoxA). After the successful placement of the model using Phaser (*36*), the FoxA model was completed using a combination of phenix.autobuild (*37*) and manual building in Coot (*38*). Apo FoxA was refined using phenix.refine and REFMAC5 (*39*) (*40*). For the FoxA-FoaB complex, refinement was performed initially using phenix.refine, followed by TLS and jelly-body refinement in REFMAC5 (*40*). For modelling the FoxA-FoaB-TonB_Ct_ complex, apo FoxA was used as a search model in Phaser. Once the MR solution was identified, clear density corresponding to the TonB_Ct_ and the N-terminal signalling domain became evident and they were manually built into the electron density using Coot. Refinement was carried out initially using phenix.refine at the early stages of model building. Once all the backbone poly-Ala stretches were built, Buster-TNT (*41*) was used for all the subsequent refinement procedures. The final models correspond to residues 44-820 of FoxA with 119-124 and 138-155 being disordered. The TonB_Ct_ model comprises residues 251-340. All data collection and refinement statistics are summarized in Table I.

**Table 1:** Data collection and refinement statistics.

|  | Apo FoxA<br>(pdb: 6I98) | FoxA-ferrioxamine B<br>(pdb: 6I96) | FoxA-ferrioxamine B-TonB<sub>ct</sub> complex<br>(pdb: 6I97) |
| --- | --- | --- | --- |
| **Data collection** |  |  |  |
| **Beamline** | PETRA III, P13 | BESSY 14.1 | PETRA III, P13 |
| Space group | P6<sub>3</sub>22 | P3<sub>2</sub>21 | P2<sub>1</sub>2<sub>1</sub>2<sub>1</sub> |
| Cell dimensions |  |  |  |
| *a*, *b*, *c* (Å) | 174.6, 174.6, 180.2 | 94.9, 94.9, 177.6 | 163.6, 174.4, 214.1 |
| $\alpha$, $\beta$, $\gamma$ (°) | 90, 90, 120 | 90, 90, 120 | 90, 90, 90 |
| Resolution (Å) | 48.53-2.80 (2.91-2.80) | 82.18-1.85 (1.88-1.85) | 48.76-3.35 (3.41-3.35) |
| *R*<sub>merge</sub> | 0.206 (1.56) | 0.15 (1.29) | 0.286 (1.8) |
| *R*<sub>meas</sub> | 0.226 (1.72) | 0.16 (1.45) | 0.34 (2.14) |
| *I* / σ*I* | 10 (1.6) | 11.2 (1.0) | 5.6 (1.0) |
| *CC*<sub>1/2</sub> | 0.99 (0.55) | 0.99 (0.28) | 0.99 (0.389) |
| Completeness (%) | 100 (99.9) | 99.2 (92.6) | 100 (100) |
| Redundancy | 10.7 (10.6) | 17 (4.7) | 6.7 (6.5) |
| **Refinement** |  |  |  |
| Resolution (Å) | 2.8 | 1.85 | 3.35 |
| No. reflections | 38331 (2769) | 75099 (5174) | 84137 (6117) |
| *R*<sub>work</sub> / *R*<sub>free</sub> | 0.21/0.26 | 0.18/0.22 | 0.23/0.26 |
| No. atoms | 5527 | 5945 | 13437 |
| Protein | 5325 | 5325 | 13357 |
| Ligand/ion | 191 | 363 | 80 |
| Water | 14 | 257 |  |
| *B*-factors |  |  |  |
| Protein | 50.2 | 49.7 | 68.1 |
| Ligand/ion | 87.2 | 81.8 | 79.3 |
| Water | 43.4 | 47.8 |  |
| R.m.s. deviations |  |  |  |
| Bond lengths (Å) | 0.02 | 0.027 | 0.02 |
| Bond angles (°) | 2.22 | 3.18 | 2.52 |

\*Values in parentheses are for highest-resolution shell.

**Table 2:** Summary of all the thermodynamic parameters determined by ITC.

| **Interaction studied** | **K<sub>d</sub>** | **ΔH (kcal mol<sup>-1</sup>)** | **TΔS (kcal mol<sup>-1</sup>)** |
| --- | --- | --- | --- |
| FoxA with TonB<sub>Ct</sub> | 107 ± 30 nM | -11.01 ± 0.3 | -1.47 |
| FoxA-foaB with TonB<sub>Ct</sub> | 6 ± 4.5 nM | -19.16 ± 0.45 | -7.86 |
| FoxA with foaB | 180 ± 140 nM | -5.07 ± 0.45 | 4.2 |
| FoxA-TonB<sub>Ct</sub> with foaB | 420 ± 300 nM | -9.37 ± 0.75 | -0.77 |

### Isothermal titration calorimetry (ITC)

FoxA was incorporated into MSP1D1 nanodiscs. Briefly, FoxA was mixed with purified MSP1D1 and POPC lipids in 1:2:70 ratio and biobeads were added to initiate the nanodisc assembly. After approximately 4-5 hours, the mixture was concentrated and purified by gel size exclusion chromatography using a Superdex S200 10/300 column. All proteins were extensively dialysed against 20 mM HEPES, 150 mM NaCl, pH 7.5 overnight. 15 μM FoxA or FoxA-FoaB incorporated into MSP1D1 nanodiscs were loaded into the ITC cell, 150 μM TonB_Ct_ was placed in the syringe. All binding reactions were measured at 26 °C. TonB_Ct_ was also titrated against empty nanodiscs as a control. Heat of dilution was obtained by titrating TonB_Ct_ into the dialysis buffer and subtracted from all subsequent measurements. Heats of binding for all the reactions were integrated using Microcal Origin software and all the data were fitted to a single-site binding model.

### Tryptophan fluorescence quenching experiments

All fluorescence measurements were performed using a Cary Eclipse fluorescence spectrometer. FoxA purified in nonyl glucopyranoside (NG) was diluted to 100 nM in a 3 ml quartz cuvette. Trp fluorescence was excited at 280 nm and emission spectra were recorded from 310-420 nm. Ferrioxamine B, diluted in the same buffer as FoxA, was titrated into the cuvette until saturation in fluorescence quenching was reached. Control experiments with buffer were performed to account for dilution effects on Trp fluorescence. Buffer conditions were 20 mM Tris pH 7.5, 200 mM NaCl, 0.4% NG. Curves were plotted and analysed in GraphPad Prism 7. The binding curve was fitted to a single-site binding model.

## Acknowledgements

We are grateful to the staff at beamlines P13 and P14 (EMBL, Hamburg), BL14.1 (BESSY, Berlin) and ID30B (ESRF, Grenoble) and thank members of the Tidow lab for helpful discussions. We acknowledge access to the Sample Preparation and Characterization (SPC) Facility of EMBL. This research was funded by the excellence cluster ‘The Hamburg Centre for Ultrafast Imaging - Structure, Dynamics and Control of Matter at the Atomic Scale) of the Deutsche Forschungsgemeinschaft (DFG EXC 1074).

## Author Contributions

Investigation, I.J. and K.V.; Writing, I.J. and H.T.; Funding Acquisition & Supervision, H.T.

## Notes

The authors declare no competing financial interest.

## Footnotes

ASU: asymmetric unit
EPR: electron paramagnetic resonance
FoaB: ferrioxamine B
MSP1D1: membrane scaffold protein variant D1
IM: inner membrane
IPTG: isopropyl β-D-1-thiogalactopyranoside
ITC: isothermal titration calorimetry
ND: nanodisc
NG: nonyl glucopyranoside
Ni-NTA: Ni-nitrilotriacetic acid
OM: outer membrane
OG: octyl glucopyranoside
POPC: 1-palmitoyl-2-oleoyl-glycero-3-phosphocholine
TBDT: TonB-dependent transporter
TEV: tobacco etch virus.

## Data Availability

Structural coordinates and structural factors have been deposited in the RCSB Protein Data Bank under accession numbers 6I96, 6I97, and 6I98 (see Table 1, Suppl. Info.). All other relevant data generated during and/or analyzed during the current study are available from the corresponding author on reasonable request.

---

**Supplementary Figure 1.**
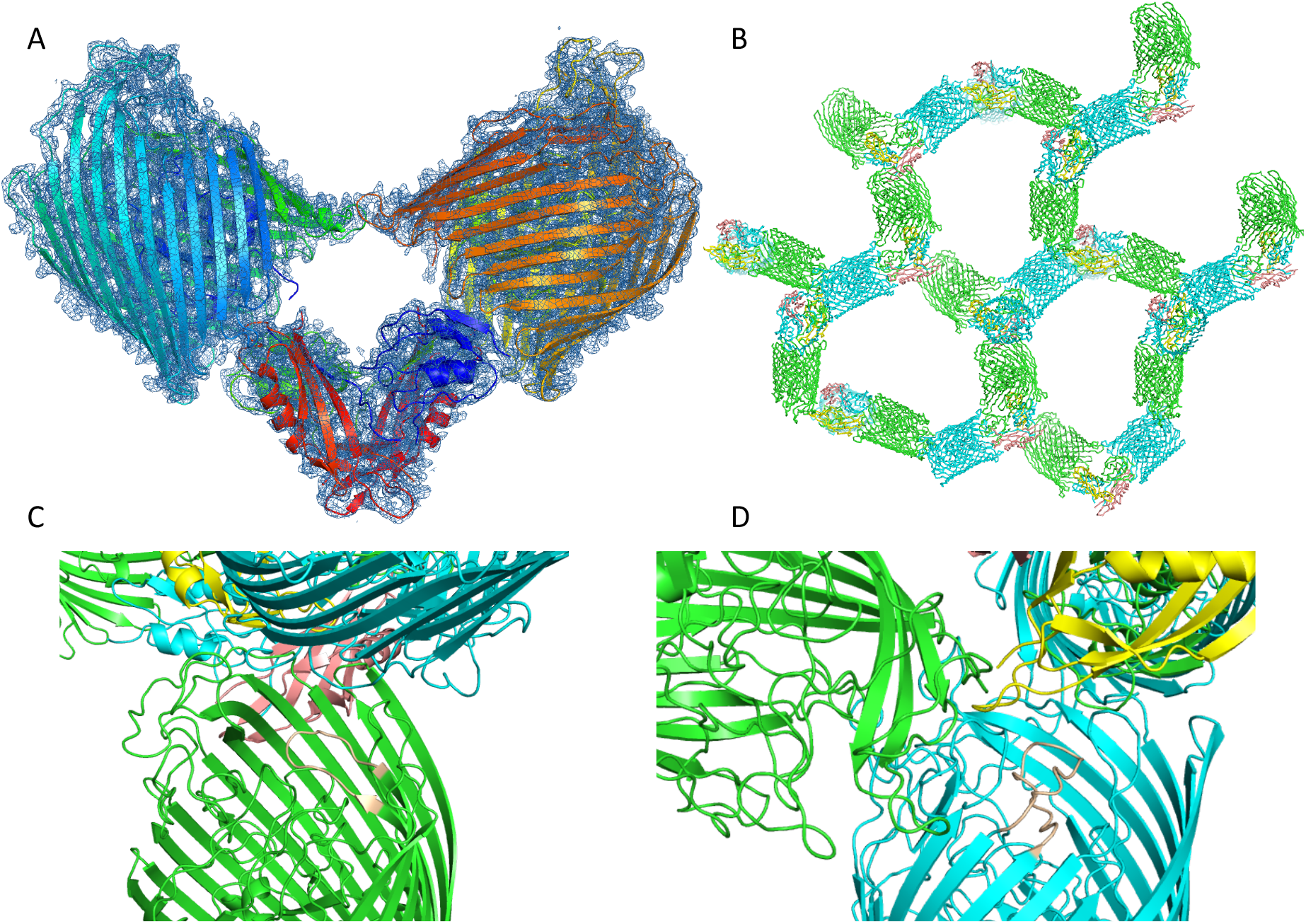
Crystal packing of the FoxA-ferrioxamine B-TonB_Ct_ complex. A) The contents of the asymmetric unit of the complex with electron density contoured at 1 σ. B) Crystal packing of the complex. C) and D) show examples of crystal packing between the FoxA complexes showing little effect of crystal packing on loop conformations between the two monomers of FoxA and the neighbouring molecules.

**Supplementary Figure 2.**
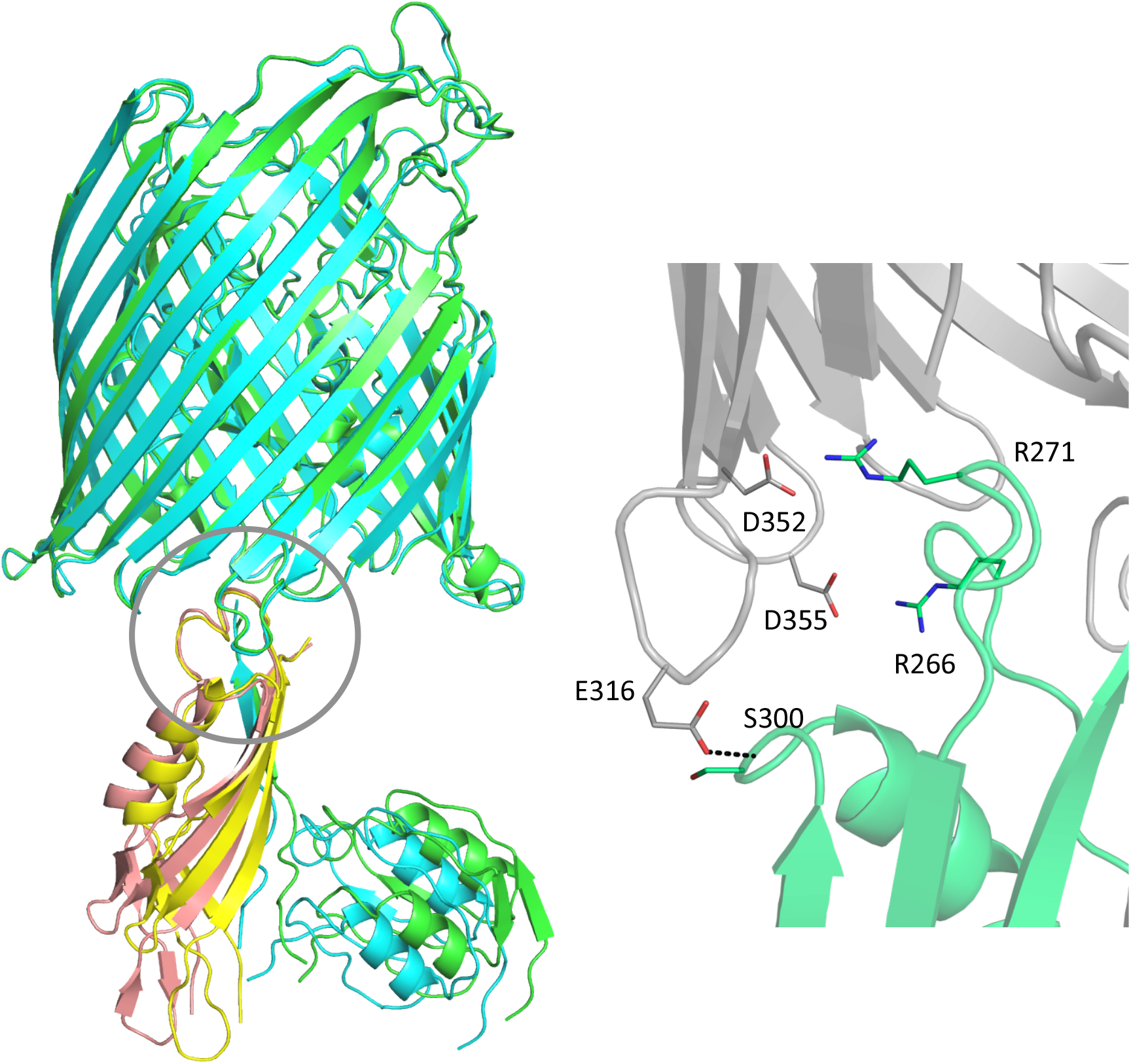
Flexibility in the FoxA-TonB complex. Overlay of the two individual complexes found in the asymmetric unit reveals that the TonB fragment and the signalling domain along with the TonB box are shifted by roughly 9° in the distal part of the complex. The protein region closest to the membrane and the barrel lumen exhibit limited movement, highlighting the importance of the secondary tethering site in FoxA. Secondary contacts between FoxA (grey) and TonB_Ct_ (green) are mediated through electrostatic interactions and hydrogen bonds.

**Supplementary Figure 3.**
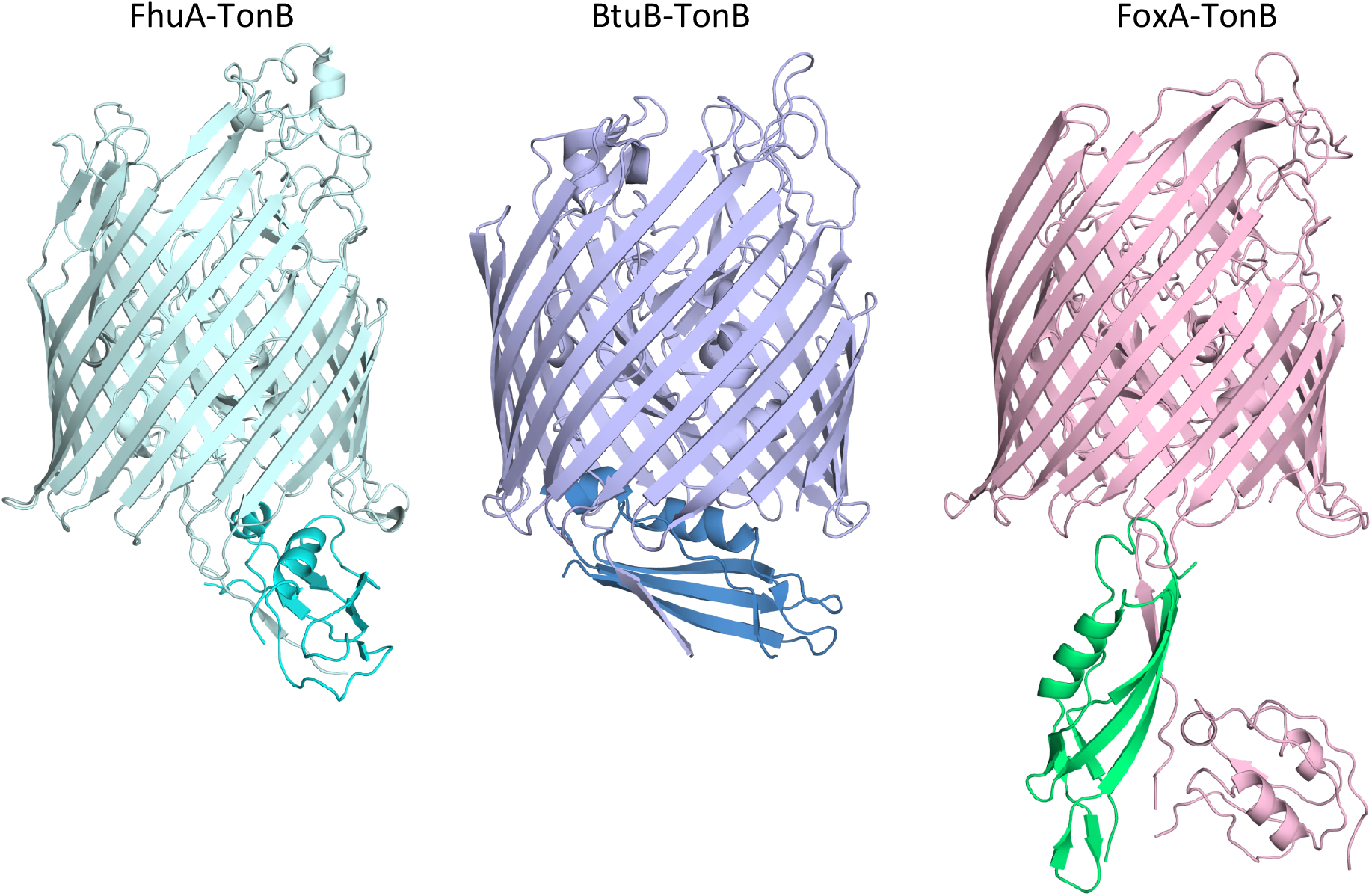
Comparison of different TBDT-TonB complex structures reveals distinct mechanisms of TonB capture and positioning. Shown are the structures of FhuA-TonB (PDBID: 2GRX) and BtuB-TonB (PDBID: 2GSK).

**Supplementary Figure 4.**
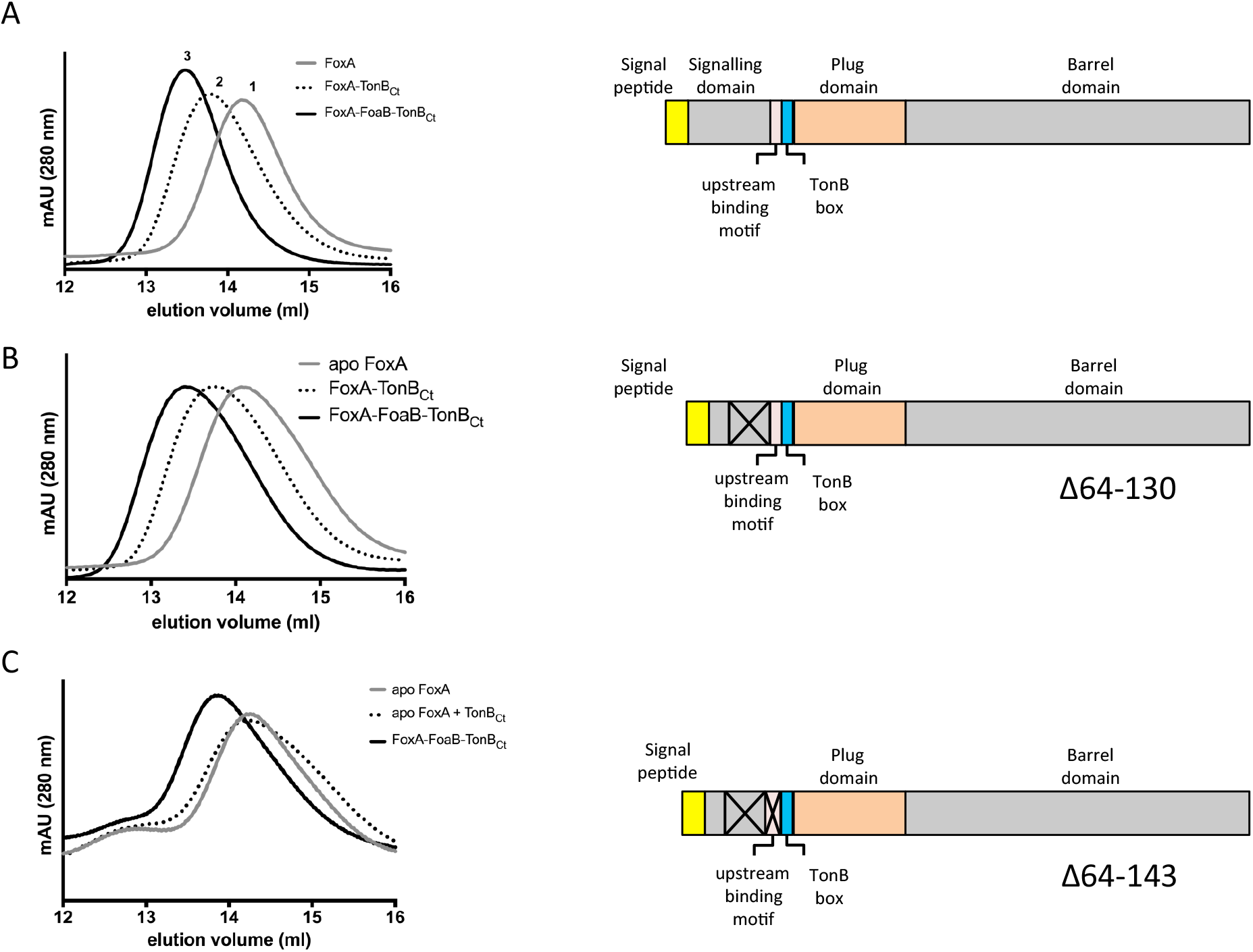
Delineation of the constitutive TonB-binding motif in FoxA using analytical size-exclusion chromatograpgy. A) Addition of TonB_Ct_ (100 μM) results in the earlier elution of the main peak when mixed with purified apo FoxA (10-20 μM). In the presence of ferrioxamine B (0.5 mM) the main peak shifts to even earlier elution volumes. B) Deletion of residues 64-130 in FoxA does not affect the elution profiles of the complexes with and without ferrioxamine B. C) Truncation of residues 64-143 from the full-length FoxA abrogates constitutive mode of binding between the receptor and TonB_Ct_, however the proteins can still associate in the presence of ferrioxamine B.

**Supplementary Figure 5.**
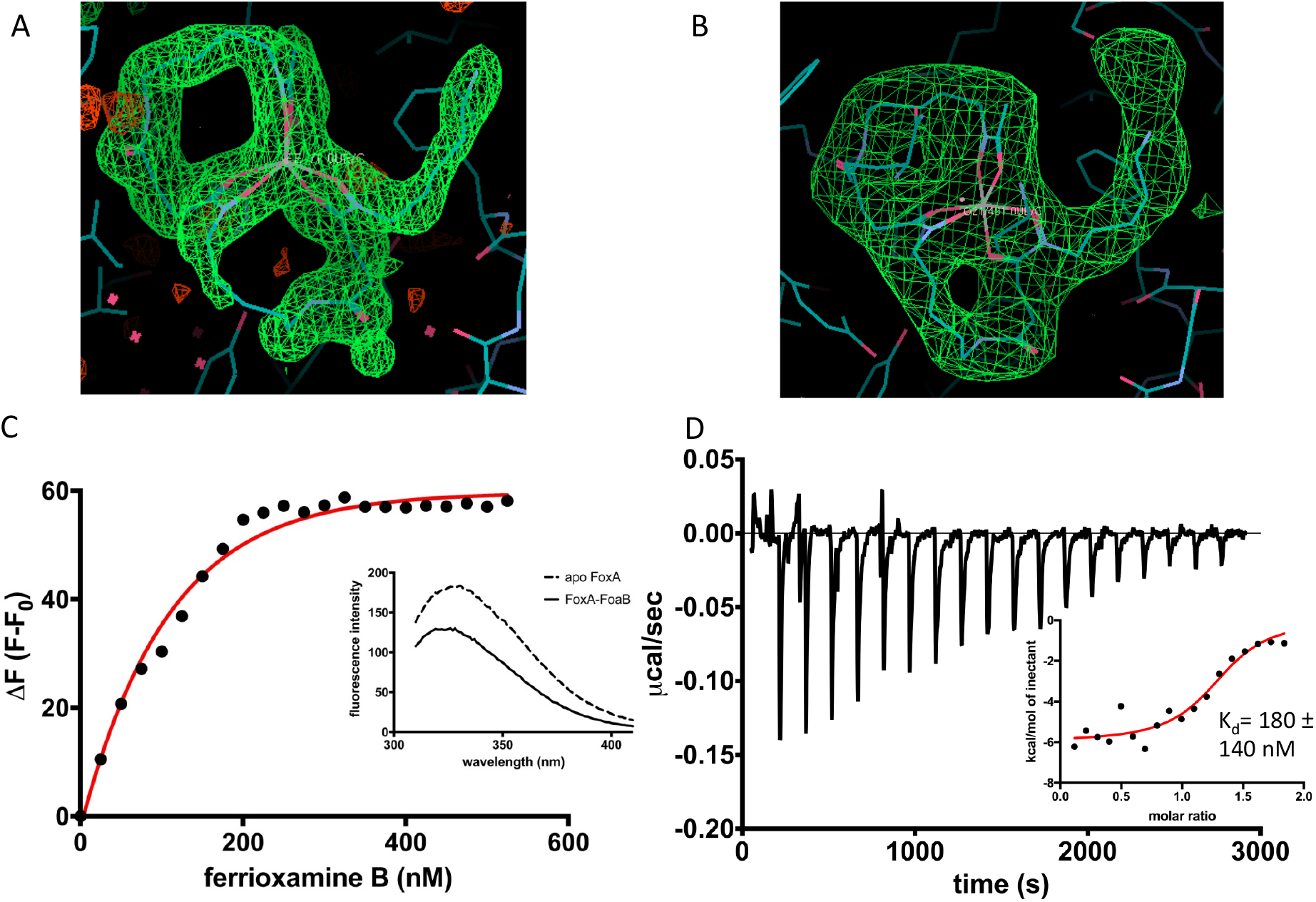
Characterization of ferrioxamine B binding to FoxA. A) Polder omit maps of ferrioxamine B in the ferrioxamine B-bound crystal structure of FoxA contoured at 3 σ. B) Polder omit map of ferrioxamine B in the ternary complex contoured at 7.2 σ. C) Titration of ferrioxamine B into purified FoxA (100 nM protein) leads to quenching of tryptophan fluorescence. The difference in fluorescence is plotted (black circles) and the data are fitted to a single-site binding model (red curve). Inset: fluorescence spectra of 100 nM FoxA purified in nonyl glucopyranoside in the absence and in the presence of saturating amounts of ferrioxamine B (500 nM). D) ITC experiment measuring ferrioxamine B titration (250 μM) into 15 μM FoxA-ND. Inset shows the integrated heats fitted to a single-site binding model with the calculated K_d_ of 180 nM.

**Supplementary Figure 6.**
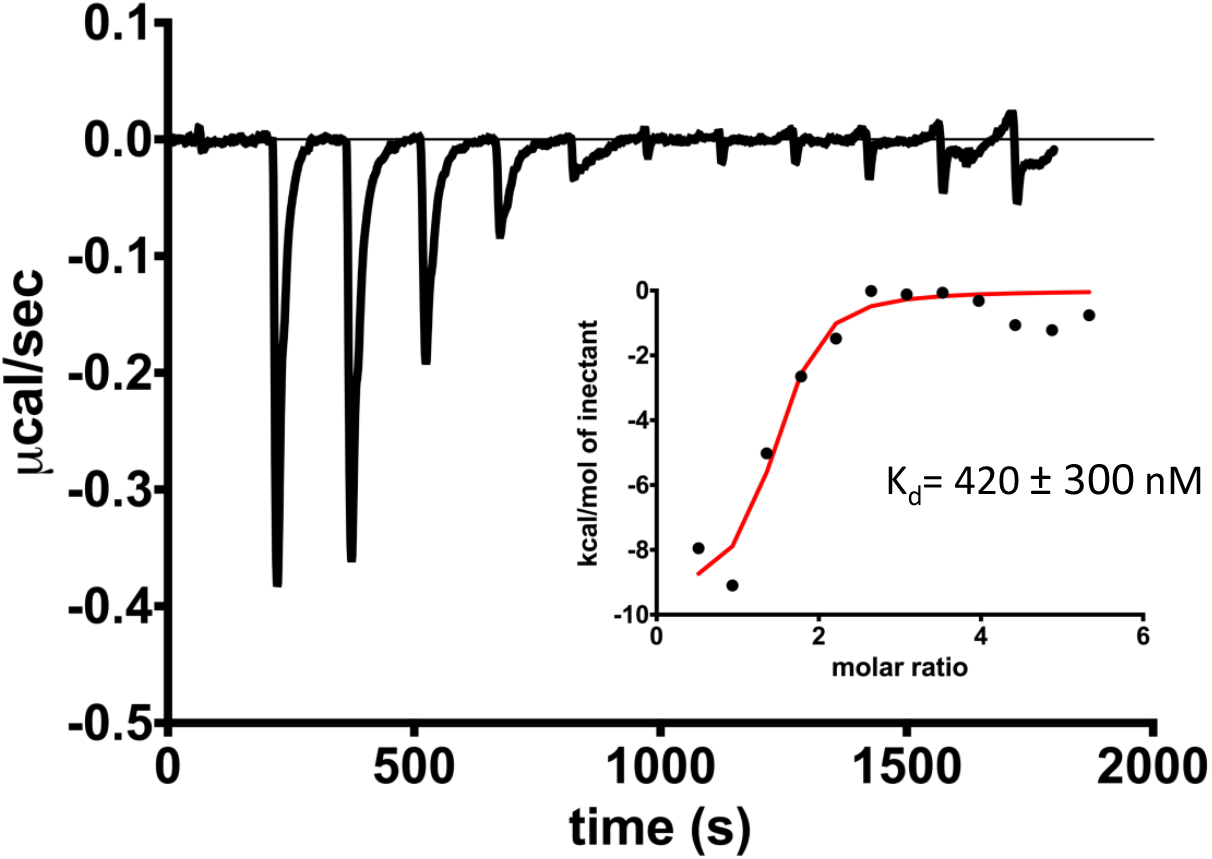
ITC measurement titrating 500 μM ferrioxamine B into 15-20 μM of pre-assembled FoxA-TonB_Ct_ in nanodiscs shows that constitutive binding of TonB_Ct_ does not lead to the closure of extracellular loops in apo FoxA.

**Supplementary Movie**

Visualizing the conformational changes occurring in FoxA in response to ferrioxamine B and TonB_Ct_ binding. We observe the closure of extracellular loops 7 and 8 as a prerequisite for translocation of ferrioxamine B. At the periplasmic side, TonB box is expelled from the plug domain in order to make contacts with the TonB molecule.

